# Determining the association between regionalisation of cortical morphology and cognition in 10,145 children

**DOI:** 10.1101/816025

**Authors:** C E Palmer, W Zhao, R Loughnan, J Zou, C C Fan, W K Thompson, T L Jernigan, A M Dale

## Abstract

Individuals undergo protracted changes in cortical morphology during childhood and adolescence, coinciding with cognitive development. Studies quantifying the association between brain structure and cognition do not always assess regional cortical morphology relative to global brain measures and typically rely on mass univariate statistics or ROI-based analyses. After controlling for global brain measures, it is possible to detect a residual regionalisation pattern indicating the size or thickness of different regions relative to the total cortical surface area or mean thickness. Individual variability in regionalisation may be important for understanding and predicting between subject variability in cognitive performance. Here we sought to determine whether the relative configuration of cortical architecture across the whole cortex was associated with cognition using a novel multivariate omnibus statistical test (MOSTest) in 10,145 children aged 9-10 years from the Adolescent Brain and Cognitive Development (ABCD) Study. MOSTest is better powered to detect associations that are widely distributed across the cortex compared to methods that assume sparse associations. We then quantified the magnitude of the association between vertex-wise cortical morphology and cognitive performance using a linear weighted sum across vertices, based on the estimated vertex-wise effect sizes. We show that the relative pattern of cortical architecture, after removing the effects of global brain measures, predicted unique variance associated with cognition across different imaging modalities and cognitive domains.

**SIGNIFICANCE STATEMENT:** This paper demonstrates a significant advance in our understanding of the relationship between cortical morphology and individual variability in cognition. There is increasing evidence that brain-behaviour associations are distributed across the cortex. Using the unprecedented sample from the Adolescent Brain and Cognitive Development (ABCD) study and a novel application of a multivariate statistical approach (MOSTest), we have discovered specific distributed regionalization patterns across the cortex associated with cognition across multiple cognitive domains. This furthers our understanding of the relationship between brain structure and cognition, namely that these associations are not sparse and localized as assumed with traditional neuroimaging analyses. This multivariate method is extremely versatile and can be used in several different applications.

## INTRODUCTION

During childhood and adolescence there are protracted changes in cortical surface area (which peaks at 9 to 10 years of age) and genetically distinct monotonic decreases in cortical thickness (1–6) that coincide with rapid cognitive development. However, the relationship between these distinct neuroanatomical trajectories and cognition is not well understood in late childhood. This is thought to be in part due to methodological differences in the estimation of cortical thickness during image pre-processing and differences in the exact morphometry measures analysed (3, 6). Many studies have looked at regional associations between brain structure and cognition; however, these do not always partition out the variance associated with modality-specific global brain measures. Regional differences in cortical morphology relative to total cortical surface area (CSA) or mean cortical thickness (CTH) appear to be important for predicting behaviour (7–11). This suggests that individual differences in the regionalisation of the cortex (relative to global brain measures) may be important for understanding individual variability in cognition. However, previous studies have been underpowered to detect significant effects of relative cortical configuration across the whole cortex particularly when using univariate statistical methods with stringent control for multiple comparisons (11). Indeed, we know that during embryonic development, regionalisation of areal expansion and apparent cortical thickness occurs via the graded expression of transcription factors across the cortical plate (12, 13). Individual variability in this patterning could therefore result in subtle changes to the whole configuration of the cortex that may impact cognition.

Brain-behaviour relationships likely represent weak diffuse effects across the brain, which are difficult to detect with small sample sizes. A recent study has highlighted how stringent thresholding procedures used in neuroimaging can produce variability among study replications and mask small, but true effects (14). From genome-wide association studies (GWAS) it has been discovered that complex behavioural traits are highly polygenic and the additive effect of many small effects across the genome is more predictive of behavioural phenotypes than effects of genome-wide significant single nucleotide polymorphisms (SNPs) chosen with bonferroni thresholding procedures (15). The same may be true of whole-brain associations. We have recently demonstrated that we can explain more variability in behaviour by taking into account effect sizes across the whole cortex using novel analysis methods adapted from genetics research (16). A *polyvertex* score (PVS), like the *polygenic risk* score (PRS), allows us to aggregate vertex-wise (or voxel-wise) effects across the cortex to predict behaviour taking into account the correlation structure among vertices. Based on this previous research and our understanding of how regionalisation occurs during development, we hypothesise that the association between cortical morphology and cognition is more likely to be widely distributed across the cortex.

In the current study, we hypothesised that the relative configuration of brain morphology (controlling for global measures) across the whole cortex would be associated with individual differences in cognitive task performance. To test this we used a multivariate omnibus statistical test (MOSTest) for determining associations between vertex-wise imaging data and behaviour. The MOSTest aggregates effects across the cortex and is therefore well-powered to detect distributed associations. This is in contrast to a commonly-used neuroimaging omnibus test based on the most significant vertex or voxel, which is better powered for sparse, localised associations. This is a particularly pertinent statistical approach given the graded nature of the biology underlying the regionalisation of the cortex in development. Van der Meer et al. recently validated this method in a different context and demonstrated the increased power of the MOSTest for detecting genome-wide associations with regional imaging phenotypes in UK Biobank (van der Meer et al., 2019). Here we demonstrate an application of the MOSTest for improving the discoverability of vertex-wise brain-behaviour associations in the Adolescent Brain and Cognitive Development (ABCD) study.

The large-scale ABCD study uses neuroimaging, genetics and a multi-dimensional battery of behavioural assessments to investigate the role of biological, environmental, and behavioural factors on brain, cognitive, and social/emotional development. The ABCD NIMH Data Archive (NDA) Release 2.0.1 includes baseline assessments for 11,878 nine and ten year old children, which allows us to generate a comprehensive cross-sectional analysis of the relationship between brain structure and cognition. Using the complete baseline data, we used the MOSTest to assess whether there was an association between performance on the thirteen cognitive tasks in the ABCD neurocognitive battery and the regionalisation of CSA (controlling for total CSA) and CTH (controlling for mean CTH) using the MOSTest. We then quantified the magnitude of these relationships using a PVS. We hypothesised that individual variability in the relative regionalisation of cortical morphology would explain cognitive task performance above and beyond that predicted by global brain measures.

## RESULTS

### Determining the association between relative cortical morphology and cognition

In order to test the association between the regionalisation of cortical morphology across the whole cortical surface and cognition in this study, we employed the MOSTest for each neurocognitive assessment and morphology measure.

#### Relative CSA

For each cognitive task, the multivariate statistic, *χ*^2^_*MOST*_, was calculated for each permutation (n=10,000) for statistical inference. In figure 1, we have plotted the distribution of *χ*^2^_*MOST*_ across permutations and superimposed the observed (non-permuted) statistic (green) to demonstrate the likelihood of observing the non-permuted statistic by chance. The Bonferroni corrected alpha threshold, based on the number of cognitive tests analysed (0.05/13=0.0038), is overlaid (red) on each plot in order to demonstrate the significance of the permutation test. The MOSTest showed a significant association between relative cortical configuration of CSA (controlling for total CSA) and cognition for all of the cognitive performance measures. We also conducted the same permutation test using the most significant vertex (*P*_*MIN*_) as the test statistic for comparison. The results are shown in supplementary figure 1. Here we can see that using the same threshold for significance the flanker task, pattern speed, picture sequencing, LMT and RAVLT did not show significant associations.

**Figure 1.**
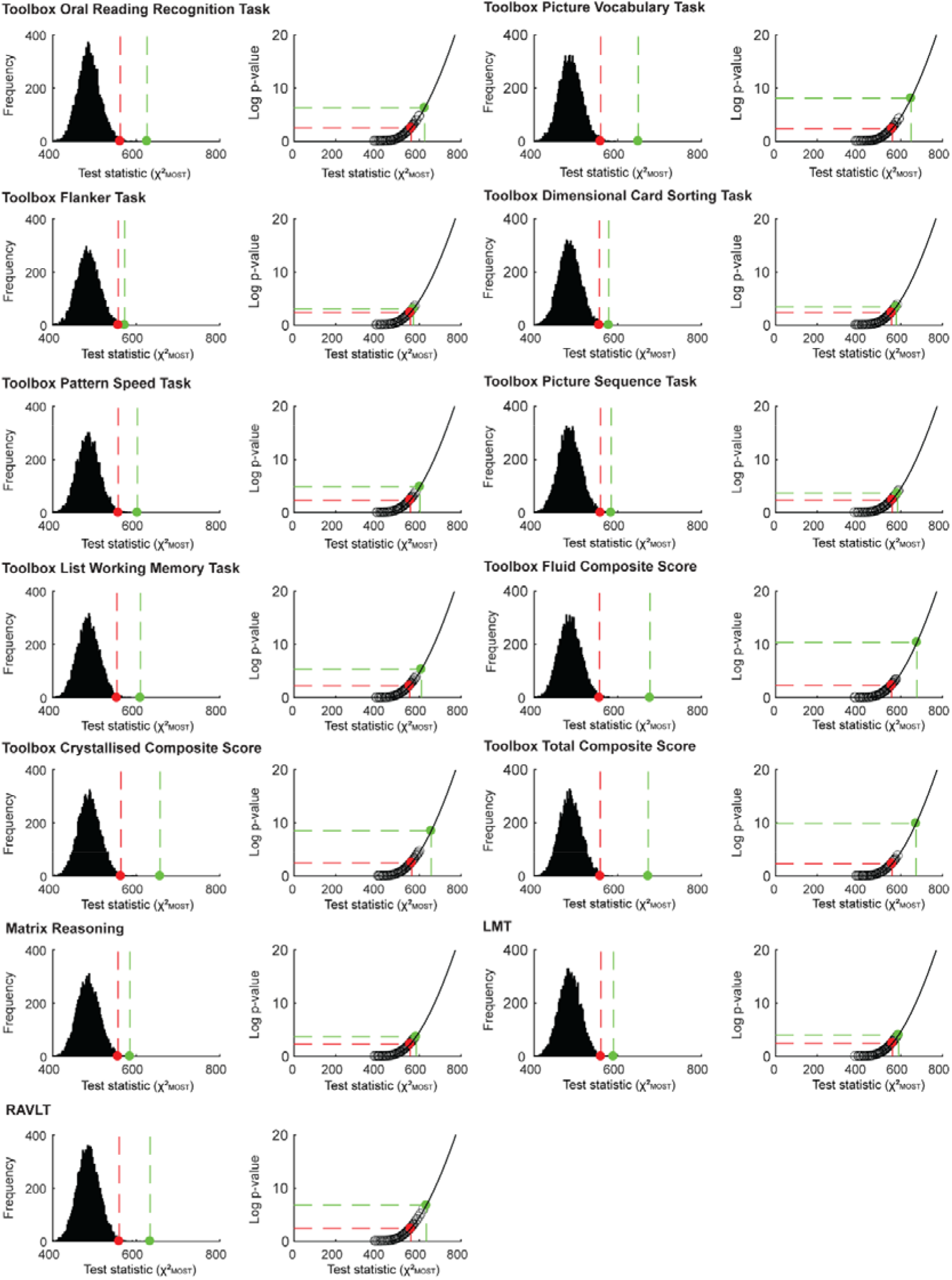
Multivariate omnibus permutation test for each cognitive task and relative CSA. The multivariate test statistic (*χ*^2^_*MOST*_) was calculated as the Mahalanobis norm of the decorrelated mass univariate z statistics for the association between relative CSA and each of the cognitive tasks. All of the left side plots show the permuted null distribution of this statistic for each cognitive task. The observed, non-permuted test statistic is shown in green and the Bonferroni corrected alpha level in red. All of the right side plots show the likelihood values for the permuted observations after fitting a gamma cumulative distribution function to the tail of the null distribution. This can be used to approximate a theoretical p-value for the observed statistic.

#### Relative CTH

In figure 2, we show the permuted and non-permuted statistics for the association between cognitive performance and the relative cortical configuration of CTH (controlling for mean CTH). The relative cortical configuration of CTH was significantly associated with all of the cognitive measures at the Bonferroni corrected alpha level (p<0.0038) according to the MOSTest. The same permutation test was carried out using the most significant vertex, *P*_*MIN*_, and the results are shown in supplementary figure 2. According to this analysis, for comparison, the following measures would not have been considered to show a significant association with relative CTH: picture vocabulary, flanker, pattern speed, LMT and RAVLT. Controlling for mean CTH does not control for brain size in the same way as controlling for total CSA, therefore in order to ensure these results were not driven by total CSA we ran the MOSTest again for relative CTH including total CSA as an additional covariate. Relative CTH was still significantly associated with all of the cognitive tasks (see Supplementary Table 3).

**Figure 2.**
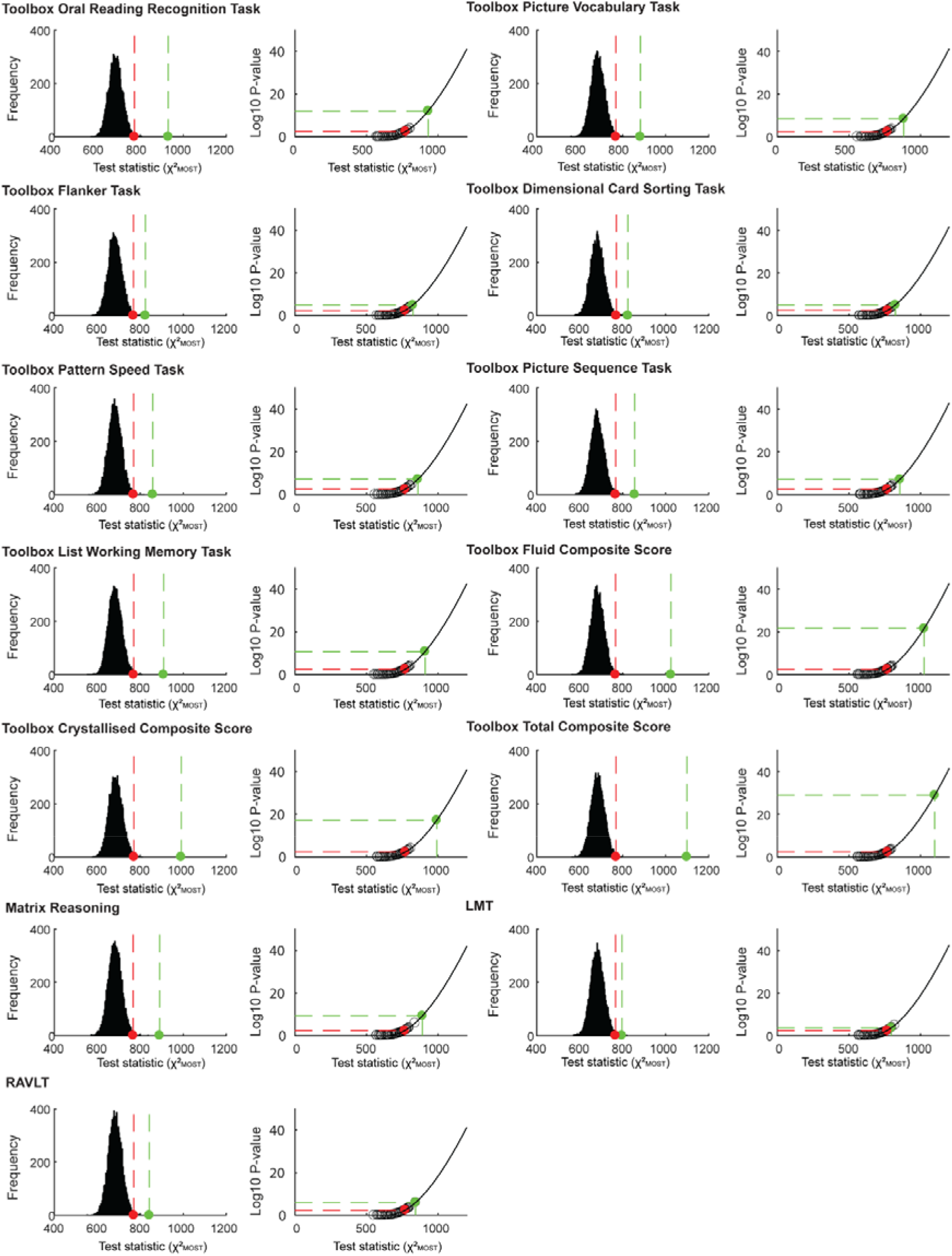
Multivariate omnibus permutation test for each cognitive task and relative CTH. The multivariate test statistic (*χ*^2^_*MOST*_) was calculated as the Mahalanobis norm of the decorrelated mass univariate z statistics for the association between relative CTH and each of the cognitive tasks. All of the left side plots show the permuted null distribution of this statistic for each cognitive task. The observed, non-permuted test statistic is shown in green and the Bonferroni corrected alpha level in red. All of the right side plots show the likelihood values for the permuted observations after fitting a gamma cumulative distribution function to the tail of the null distribution. This can be used to approximate a theoretical p-value for the observed statistic.

Across both relative CSA and CTH, the p-values for the MOSTest associations were magnitudes smaller than for the min-p associations (Supplementary Table 3). We used empirical simulations to compare the sensitivity for detection of the MOSTest and the standard approach (see Methods for details). The MOSTest showed increased sensitivity for detection across multiple random samples at different sample sizes compared to the standard approach, whilst, importantly, maintaining a false positive rate below 5% (supplementary figure 6). The increased power of the MOSTest is likely due to the distributed nature of the associations between cognition and relative cortical morphology. This can clearly be seen in the surface maps of the unthresholded mass univariate effect sizes for relative CSA and CTH predicting the total composite score from the NIH Toolbox (Figure 3).

**Figure 3.**
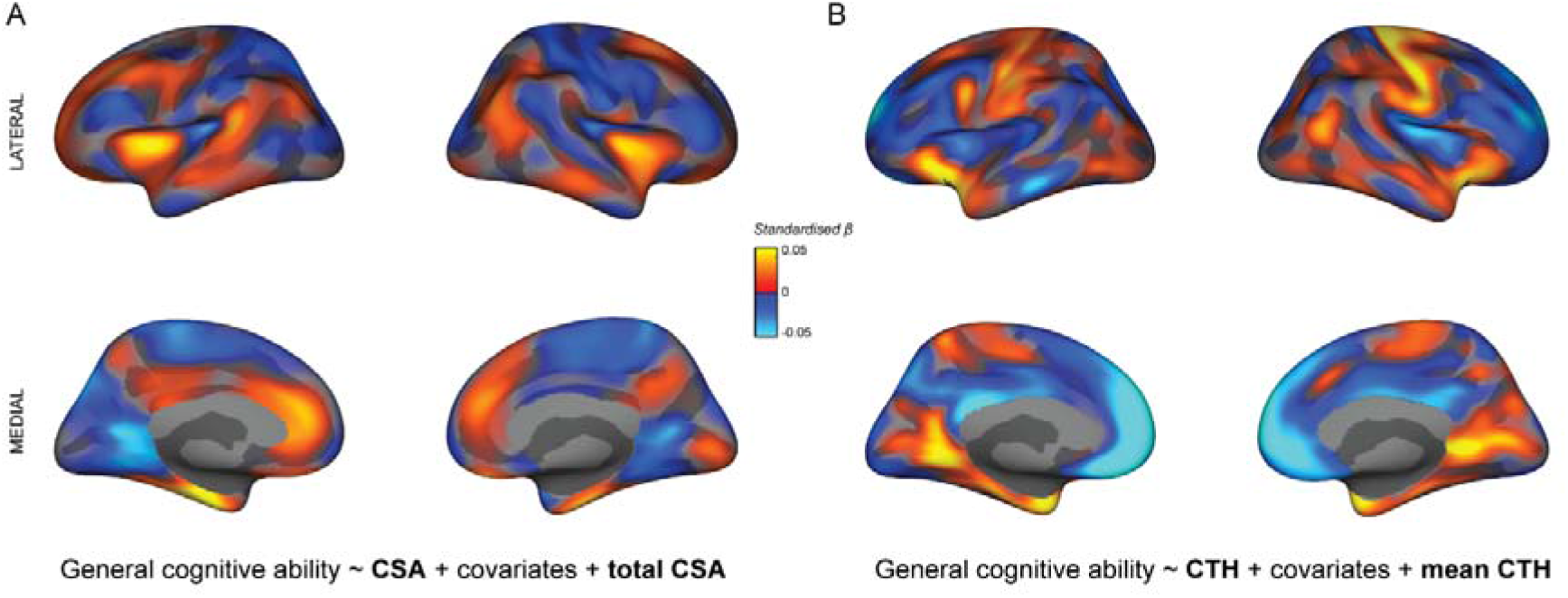
Surface maps showing the mass univariate standardised beta coefficients for the association between the NIH Toolbox Total Composite score and (A) relative CSA and (B) relative CTH. These unthresholded effect size maps show the continuously distributed associations across the cortical surface. These distributed patterns of association support the need to use a multivariate statistic that incorporates this entire pattern to determine the relationship between relative cortical morphology and cognition.

### Quantifying the association between brain structure and cognition

Having established there was a significant association between relative cortical configuration and cognition across tasks, we aimed to quantify the unique proportion of individual variability in each behaviour predicted by the relative cortical configuration. To do this we used the PVS_B_ in order to provide a conservative, out-of-sample estimate of the variance in each behaviour predicted by the relative vertex-wise imaging data. Each PVS_B_ was orthogonal to the modality specific global imaging measures (total CSA and mean CTH respectively) as all imaging data were pre-residualised using these measures (and the additional covariates of no interest) prior to calculating the PVS_B_. We then used a linear model to determine the unique contribution of each PVS_B_ (relative CSA and relative CTH) to cognitive task performance and compared to this to the independent contribution of the global imaging measures. Figure 4 shows the variance (%R^2^) in each cognitive task performance explained by total CSA (dark blue), relative CSA (light blue), mean CTH (dark pink) and relative CTH (light pink).

**Figure 4.**
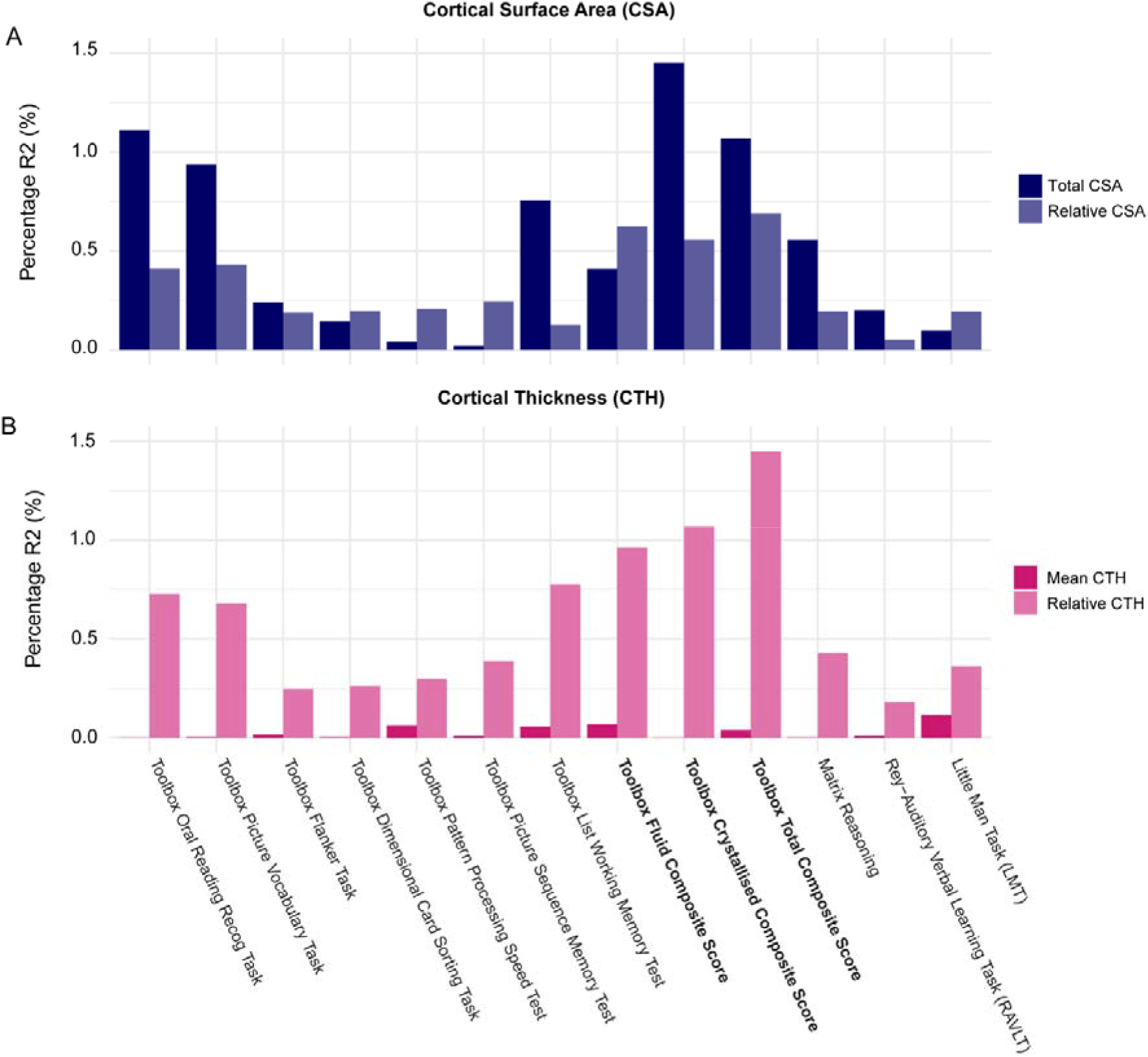
Percent behavioural variance explained across cognitive tasks by CSA and CTH. Variance explained was calculated by correlating each PVS with the observed behaviour for each cognitive task to generate a %R^2^ value. Each PVS was generated by aggregating the Bayesian estimated effect sizes across the whole cortex from the association between the imaging data and behaviour. The composite scores, which represent averages of single task performance, are highlighted in bold. The percent variance explained for each cognitive measure was determined for the following imaging phenotypes: CSA (dark blue), relative CSA (controlling for total CSA; light blue), CTH (dark pink), relative CTH (controlling for mean CTH; light pink).

The relative configuration of CSA and CTH explained a greater proportion of unique variance in the composite scores (relative CSA: 0.96-1.31%R^2^; relative CTH: 1.06-1.38%R^2^) compared to the single task measures. For the single measures, more variability in cognitive performance was explained in the crystallised measures (reading and vocabulary) and list working memory (relative CSA: 0.78-0.91%R^2^; relative CTH: 0.60-0.80%R^2^) compared to the individual fluid measures (relative CSA: 0.11-0.34%R^2^; relative CTH: 0.27-0.38%R^2^).

Total CSA predicted more variance in cognitive performance compared to relative CSA for the crystallised measures (reading recognition, picture vocabulary, crystallised composite score and the total composite score); however, this relationship was not always true for the other cognitive measures. Indeed, for the fluid composite score, total and relative CSA predicted a similar proportion of variance. This suggests that brain size is more predictive of crystallised compared to fluid cognitive measures. Mean CTH was not predictive of cognitive performance across any domains, however, relative CTH was predictive across cognitive domains and appeared more predictive than relative CSA. However, this may be misleading as mean CTH does not control for brain size in the same way as total CSA. Therefore, we computed the unique variance in cognition explained by relative CTH when controlling for total CSA as well as mean CTH (supplementary figure 7). The percentage R^2^ decreased by a greater proportion for the crystallised compared to the fluid measures, which is unsurprising given that these measures show a greater total CSA effect. The variance explained by the relative cortical morphology measures was more comparable across tasks after both imaging phenotypes controlled for total CSA.

In general, the variance in behaviour explained by these morphology measures was very low; however, in this analysis we have residualised for a wide range of confounding demographic variables (race and ethnicity, household income and parental education), which removes a large portion of shared variance amongst these variables, brain structure and cognition. These estimates of R^2^ therefore provide a conservative lower bound of the contribution of cortical morphology to cognition that is not attributable to confounding variables.

## DISCUSSION

In this study, we have shown that individual differences in the regionalisation of cortical morphology were associated with cognitive performance across multiple domains in a large sample of 9 and 10 year old children. The novel multivariate statical omnibus test (MOSTest) employed here, aggregates many small effects across the cortical surface to improve our power for discovering distributed brain-behaviour associations. The MOSTest has allowed us to uncover patterns of association across the cortical surface between the regionalisation of cortical morphology and cognition that have not previously been detected. We then used a linear weighted sum of the associations (PVS_B_) to quantify the effect size of these discovered associations. The regionalisation of CSA and CTH predicted a similar magnitude of unique variance in cognition beyond that predicted by global brain measures, however the relative contribution of regionalisation and total CSA in predicting behaviour differed depending on the cognitive domain being measured. These results highlight the importance of studying the configuration of brain structure relative to global brain measures. Individual differences in these regionalisation patterns may uncover important mechanisms underlying the development of cognitive processes.

### Relative cortical configuration is important for cognition

Global brain measures have been previously associated with cognition; however, there are many examples where there are differences in global brain measures, such as brain size between men and women, which do not correspond with differences in cognition. When measuring regional associations with behaviour it is therefore essential to control for these global measures in order to uncover specific relationships between regionalisation and behaviour. Indeed, a number of studies have highlighted associations between regional differences in cortical morphology and cognition; however, they have lacked the power and multivariate methods to measure associations specifically based on the entire configuration of cortical morphology. Importantly, the regionalisation of CSA and CTH has been associated with distinct, continuous gradients of genetic influences across the cortex (18–20) that coincide with the gene expression patterns that dictate specialisation of the neocortex during embryonic development (12, 13). Individual differences in this molecular signalling could therefore lead to subtle alterations in this cortical configuration, which could lead to variability in behaviour. It is therefore important to use a statistical test that captures the graded and distributed nature of the biology of regionalisation, such as the MOSTest.

Here we have shown that performance across multiple cognitive tasks was associated with the regionalisation of cortical morphology across the whole cortex. The presence of negative and positive associations across the cortical surface suggests that it is the patterning of cortical morphology that may be important for supporting cognitive function. Indeed, the regionalisation of CSA predicted additional unique variance in cognition not explained by total CSA. This allows us to partition the variance associated with cortical morphology in order determine the relative importance of these components of CSA for cognition. This appeared to differ depending on the cognitive task, with more crystallised measures having a greater association with total CSA compared to the regionalisation of CSA. Comparatively the proportion of variance in behaviour explained by regional CTH was relatively similar for fluid and crystallised measures. Mean CTH did not predict cognitive performance across any measures; however this may be due to the developmental stage of this sample. However, additionally controlling for total CSA reduced the variance in behaviour explained by the regionalisation of CTH. Although total CSA and mean CTH are thought to be independent, there appears to be some overlap between the regionalisation of CTH and total CSA in predicting cognition.

It is important to note that the effect sizes reported between brain structure and cognition in this study are relatively small (~1% R^2^). Here we have used a very conservative, out-of-sample effect size estimate and have residualised for a number of confounding demographic variables, which are highly collinear with these measures. We are therefore providing the lower bound for the estimated effect size between regionalisation and cognition, which is not influenced by any confounding measures. Nevertheless, cognitive processing is highly complex, therefore we are unlikely to explain large proportions of individual variability using single neuroimaging modalities. Future work combining imaging measures within multi-modal predictive models will likely uncover larger associations between brain phenotypes and cognition.

### A novel multivariate vertex-wise statistical inference tool

In order to make a statistical inference about the association between cortical configuration and cognition in this study we have shown a novel application of the MOSTest for detecting vertexwise associations with behaviour. This test uses a statistic that aggregates effects of interest across the cortex; therefore, it is optimally powered to detect effects that are distributed across the cortical surface. Traditionally, neuroimaging analysis methods assume neuroimaging effects are sparse and localised, therefore adopt the most significant vertex or voxel as an omnibus test statistic. However, this can lead to reduced statistical power and an increase in Type II errors (14) particularly if the association between the brain and behaviour is continuous and distributed across the cortex. The continuous distribution of brain-behaviour associations has recently been highlighted across modalities using large samples such as the ABCD study. Indeed, a recent paper introducing the PVS, demonstrated that aggregating information across the cortex could produce greater out-of-sample prediction performance of task fMRI-behaviour associations than thresholding based on significance level (16). This mirrors findings from genetics, which demonstrate that many complex behavioural phenotypes are polygenic: genome-wide significant single nucleotide polymorphisms (SNPs) only predict a very small proportion of the variance in complex behavioural phenotypes, however collating effects across the entire genome increases the variance explained. Zhao and colleagues have shown that the same is true when integrating effects across the cortex.

In this study, the MOSTest detected a greater number of significant associations across tasks compared to using the most significant vertex as an omnibus test. According to our empirical simulations, the MOSTest had a greater sensitivity for detecting non-null vertex-wise effects compared to the standard univariate omnibus test (min-P) whilst maintaining a low false positive rate at 5% (supplementary figure 6). This demonstrates the increased power of this test for the discovery of vertex-wise associations with behaviour. Van der Meer and colleagues have applied the MOSTest to detect significant genome-wide associations with regional neuroimaging phenotypes using UK Biobank data and demonstrated a 3-fold increase in statistical power using this method compared to standard univariate analyses (17). This method improves power by capitalising on the shared variance across vertices and by negating the need to use stringent vertex-wise correction for multiple comparisons across the cortical surface; but importantly it does not inflate the type-1 error rate due to the use of a permutation test for statistical inference. In addition, the MOSTest test statistic takes into account the correlation structure of the residuals across vertices. The multivariate test statistic, the Mahalanobis norm, represents the sum of squared deviations of the vector of decorrelated mass univariate z statistics. Using this multivariate method we can readily test for associations between the brain and behaviour using information across the entire cortex with increased statistical power. Although it is clear that not all behaviours will be associated with widespread cortical activity or brain structure, these data suggest that we should not assume sparseness of neuroimaging associations.

## Conclusions

In this paper, we have shown that areal expansion of the cortex and regionalisation of apparent cortical thickness relative to global brain structure is predictive of cognitive performance across a number of different cognitive domains. Previous studies have been unable to detect significant associations between these regionalisation phenotypes and cognition due to a lack of statistical power and appropriate multivariate methods. Here we utilised the large sample size of ABCD to test for these associations and importantly applied a more statistically appropriate omnibus method for discovering brain-behaviour associations that are distributed across the cortical surface. It is essential that we used novel methods such as this to understand how individual differences in complex behaviours relate to individual differences in brain phenotypes and their development. Future work will aim to understand how these structural associations change over time with longitudinal assessments in ABCD.

## MATERIALS & METHODS

### LEAD CONTACT AND MATERIALS AVAILABILITY

Further information and requests for resources should be directed to and will be fulfilled by the Lead Contact, Clare Palmer.

### SUBJECT DETAILS

The ABCD study is a longitudinal study across 21 data acquisition sites following 11,878 children starting at 9 and 10 years old. This paper analysed the full baseline sample from release 2.0.1 (NDAR DOI: 10.15154/1504041). The ABCD Study used school-based recruitment strategies to create a population-based, demographically diverse sample, although not necessarily representative of the U.S. national population (21, 22). Due to the inclusion of a wide range of individuals across different races, ethnicities and socioeconomic backgrounds, it is important to carefully consider how to control for these potentially confounding factors and the implications of this on our effects of interest. Sex and age were also used as covariates in all analyses. Embedded within the sample is a large twin cohort and many siblings, therefore family relatedness was also controlled for as a random effect in all analyses using a restricted exchangeability permutation scheme (see *Statistical Analysis 1*).

Only participants who had complete data across all of the measures analysed were included in the neuroimaging analyses. However, an exception was made for household income due to the large number of missingness for this variable (n=1018); in order to include those participants, missing income values were imputed by taking the median income value across participants from the same testing site. The sources of missing data were as follows: incomplete across all demographic variables (n=189), incomplete across all cognitive measures (n=944), unavailable T_1_-weighted MRI scan for reasons outlined in the ABCD release notes (e.g. did not get scanned, motion artefacts) (n=339) and imaging data that was made available but did not pass the FreeSurfer QC flag (n=462). These missing data values are not mutually exclusive. The permutation testing used in this study was dependent on having multiple families with the same number of children, therefore the single family with 5 children was excluded from these analyses. This resulted in a final sample of 10,145 subjects. Supplementary Table 1 displays the names of each variable used in these analyses from data release 2.0.1. Supplementary Table 2 shows the distribution of the analysed sample stratified by the demographic covariates of no interest included in the study. All levels of these variables were represented in this subsample.

## METHOD DETAILS

### Neurocognitive assessment

#### NIH Toolbox^®^

The NIH Toolbox Cognition Battery^®^ (NTCB) is a widely used battery of cognitive tests that measures a range of different cognitive domains. All of the tasks within the NTCB were administered using an iPad with support or scoring from a research assistant where needed. Below is a brief description of each task.

The ***Toolbox Oral Reading Recognition Task***^®^ ***(TORRT)*** measured language decoding and reading. Children were asked to read aloud single letters or words presented in the center of an iPad screen. The research assistant marked pronunciations as correct or incorrect. Extensive training was given prior to administering the test battery. Item difficulty was modulated using computerised adaptive testing (CAT).

The ***Toolbox Picture Vocabulary Task***^®^ ***(TPVT)***, a variant of the Peabody Picture Vocabulary Test (PPTV), measured language and vocabulary comprehension. Four pictures were presented on an iPad screen as a word was played through the iPad speaker. The child was instructed to point to the picture, which represented the concept, idea or object name heard. CAT was implemented to control for item difficulty and avoid floor or ceiling effects.

The ***Toolbox Pattern Comparison Processing Speed Tes***^®^***t (TPCPST)*** measured processing speed. Children were shown two images and asked to determine if they were identical or different by touching the appropriate response button on the screen. This test score is the sum of the number of items completed correctly in the time given.

The ***Toolbox List Sorting Working Memory Test***^®^ ***(TLSWMT)*** measured working memory. Children heard a list of words alongside pictures of each word and were instructed to repeat the list back in order of their actual size from smallest to largest. The list started with only 2 items and a single category (e.g. animals). The number of items increased with each correct answer to a maximum of seven. The child then progressed to the next stage in which two different categories were interleaved. At this stage children were required to report the items back in size order from the first category followed by the second category. Children were always given two opportunities to repeat the list correctly before the experimenter scored the trial as incorrect.

The ***Toolbox Picture Sequence Memory Test***^®^ ***(TPSMT)*** measured episodic memory. On each trial, children were shown a series of fifteen pictures in a particular sequence. The pictures illustrated activities or events within a particular setting (e.g. going to the park). Participants were instructed to touch the pictures in the original sequence in which they were shown. The Rey-Auditory Verbal Learning Task (RAVLT) was also included in the ABCD neurocognition battery as a more comprehensive measure of episodic memory.

The ***Toolbox Flanker Task***^®^ ***(TFT)*** measured executive function, attentional and inhibitory control. This adaptation of the Eriksen Flanker task (23) captures how readily a participant is influenced by the congruency of stimuli surrounding a target. On each trial a target arrow was presented in the center of the iPad screen facing to the left or right and was flanked by two additional arrows on both sides. The surrounding arrows were either facing in the same (congruent) or different (incongruent) direction to the central target arrow. The participant was instructed to push a response button to indicate the direction of the central target arrow. Accuracy and reaction time scores were combined to produce a total score of executive attention, such that higher scores indicate a greater ability to attend to relevant information and inhibit incorrect responses.

***The Toolbox Dimensional Change Card Sort Task***^®^ ***(TDCCS)*** measured executive function and cognitive flexibility. On each trial, the participant was presented with two objects at the bottom of the iPad screen and a third object in the middle. The participant was asked to sort the third object by matching it to one of the bottom two objects based on either colour or shape. In the first block participants matched based on one dimension and in the second block they switched to the other dimension. In the final block, the sorting dimension alternated between trials pseudorandomly. The total score was calculated based on speed and accuracy.

In the current study, the uncorrected scores for each task were used for statistical analyses. Composite scores of crystallised intelligence (mean of TPVT and TORRT), fluid intelligence (mean of TPCPST, TLSWMT, TPSMT, TFT and TDCCS) and total score (mean of all tasks) were also analysed. These measures are highly correlated with ‘gold standard’ measures of intelligence in adults (24) and children (25).

#### Rey-Auditory Verbal Learning Task (RAVLT)

This task measures auditory learning, recall and recognition. Participants listened to a list of 15 unrelated words and were asked to immediately recall these after each of five learning trials. A second unrelated list was then presented and participants were asked to recall as many words as possible from the second list and then recall words again from the initial list. Following a delay of 30 minutes (during which other non-verbal tasks from the cognitive battery are administered), longer-term retention was measured using recall and recognition. This task was administered via an iPad using the Q-interactive platform of Pearson assessments (26). In the current study, the total number of items correctly recalled across the five learning trials was summed to produce a measure of auditory verbal learning.

#### Little Man Task (LMT)

This task measures visuospatial processing involving mental rotation with varying degrees of difficulty (27). A rudimentary male figure holding a briefcase in one hand was presented on an iPad screen. The figure could appear in one of four positions: right side up vs upside down and either facing the participant or with his back to the participant. The briefcase could be in either hand. Participants indicated which hand the briefcase was in using one of two buttons. Performance across the 32 trials was measured by the percentage of trials in which the child responded correctly. This was divided by the average reaction time to complete the task (in seconds) to produce a measure of efficiency of visuospatial processing. This was the dependent variable analysed in this study.

#### Matrix reasoning

Nonverbal reasoning was measured using an automated version of the Matrix Reasoning subtest from the Weschler Intelligence Test for Children-V (WISC-V; Weschler, 2014). On each trial the participant was presented with a series of visuospatial stimuli, which was incomplete. The participant was instructed to select the next stimulus in the sequence from four alternatives. There were 32 possible trials and testing ended when the participant failed three consecutive trials. The total raw score, used in the current study, was the total number of trials completed correctly.

### MRI acquisition

The ABCD MRI data were collected across 21 research sites using Siemens Prisma, GE 750 and Philips 3T scanners. Scanning protocols were harmonised across sites. The T1w acquisition (1 mm isotropic) was a 3D T_1_-weighted inversion prepared RF-spoiled gradient echo scan using prospective motion correction, when available (29, 30) (echo time = 2.88 ms, repetition time = 2500 ms, inversion time = 1060 ms, flip angle = 8°, FOV = 256×256, FOV phase = 100%, slices = 176). Only the T1w scans were analysed in this paper. Full details of all the imaging acquisition protocols used in ABCD are outlined by Casey et al (31). Scanner ID was included in all analyses to control for any differences in image acquisition across sites and scanners.

### Image pre-processing

Pre-processing of all MRI data for ABCD was conducted using in-house software at the Center for Multimodal Imaging and Genetics (CMIG) at University of California San Diego (UCSD) as outlined in Hagler et al (32). Manual quality control was performed prior to the full image pre-processing and structural scans with poor image quality as well as those that did not pass FreeSurfer QC were excluded from all analyses. Brain segmentation and cortical surface reconstruction were completed using FreeSurfer v5.3.0 (33–37).

T_1_-weighted structural images were corrected for distortions caused by gradient nonlinearities, coregistered, averaged, and rigidly resampled into alignment with an atlas brain. See previous publications for details of the surface based cortical reconstruction segmentation procedures (33, 34, 38–40). In brief, a 3D model of the cortical surface was constructed for each subject. This included segmentation of the white matter (WM), tessellation of the gray matter (GM)/WM boundary, inflation of the folded, tessellated surface, and correction of topological defects.

Measures of cortical thickness at each vertex were calculated as the shortest distance between the reconstructed GM/WM and pial surfaces (38). To calculate cortical surface area, a standardised tessellation was mapped to the native space of each subject using a spherical atlas registration, which matched the cortical folding patterns across subjects. Surface area of each point in atlas space was calculated as the area of each triangle. This generated a continuous vertex-wise measure of relative areal expansion or contraction. Cortical maps were smoothed using a Gaussian kernel of 20 mm full-width half maximum (FWHM) and mapped into standardised spherical atlas space. Vertex-wise data for all subjects for each morphometric measurement were concatenated into matrices in MATLAB R2017a and entered into general linear models for statistical analysis using custom written code.

## QUANTIFICATION AND STATISTICAL ANALYSIS

### Statistical Analysis 1: determining the association between regionalisation and cognition using the MOSTest

In order to determine whether there was a significant association between vertex-wise cortical morphology (controlling for global measures) and cognitive task performance, we employed the MOSTest. We applied a general linear model (GLM) associating a given behaviour *y* from a set of covariates *W* and the vertex-wise morphology data, *X*_*v*_

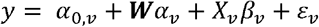

Here, *W* represents a standardized *N × m* matrix of covariates of no interest where *N* represents the number of subjects and *m* the number of covariates. *X*_*v*_ denotes the standardised *N ×* 1 vector of imaging data for the *v*th vertex. This GLM was applied univariately at each vertex v = 1, … V. Let ***β*** = (*β*_1_, …, *β*_*v*_)′ and *α* denote the *V ×* 1 and *m ×* 1 vectors of parameters of interest and no interest, respectively. Using the least-squares estimates *β*_v_ from each GLM, we computed the *V ×* 1 vector of Wald statistics **z** = (*z*, …, *z*_*v*_)′ across the whole cortex. The multivariate omnibus test statistic *χ*^2^_*MOST*_ was calculated as the estimated squared Mahalanobis norm of **z**

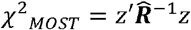

Assuming that **z** is asymptotically drawn from a multivariate normal distribution and 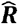 is a consistent estimator of the covariance matrix ***R***, the MOSTest statistic would have an approximate Chi-squared distribution (a special case of the gamma distribution) under the null hypothesis. However, we relax this assumption and instead compute its p-value using a hybrid permutation procedure as follows. First, the covariance matrix 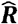 was estimated from the *V × L* matrix of permuted Wald statistics, *Z*_*perm*_, where *L* denotes the number of permutations. This was then Tikhonov regularized with penalty parameter *λ* to ensure the covariance matrix was invertible.

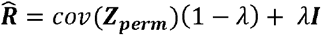

Capitalizing on the theoretical asymptotic gamma distribution of *χ*^2^_*MOST*_, we chose *λ* based on minimizing the negative log likelihood of the fitted gamma cumulative distribution function (cdf) with two free parameters to the permuted distribution of *χ*^2^_*MOST*_ (see supplementary figure 3). To correctly estimate the null distribution from permutations, subject labels were shuffled according to exchangeability blocks (EBs) defined based on the family structure within ABCD (41). Families with the same number of siblings were allowed to be shuffled as a whole and siblings within a family were allowed to be shuffled with each other. This was necessary to account for the joint distribution of the observed data due to the relatedness of the ABCD sample. Permutations that adhered to this shuffling scheme were generated using the PALM toolbox (42). Permutations were conducted using the Freedman-Lane procedure(43).

This permutation procedure was used to determine the distribution of *χ*^2^_*MOST*_ under the global null hypothesis H0. We extrapolated observed permuation p-values for *χ*^2^_*MOST*_. by again utilizing a gamma cdf fitted to the permuted null test statistics. Supplementary figures 4A & 5A show the good fit of the gamma distributions to the permuted data. We rejected H_0_ if *χ*^2^_*MOST*_ was greater than the value of *χ*^2^_*perm*_ at the critical threshold corresponding to an alpha level of 0.0038 (0.05 corrected for the 13 cognitive tests analysed).

All analyses were also conducted using the most significant vertex, *P*_*MIN*_= min (***p***), as the omnibus statistic, where ***p*** denotes the *V ×* 1 vector of p-values from each univariate vertex-wise GLM. The cumulative distribution function of this statistic follows a Sidak distribution, which is a special case of the beta distribution. Approximate p-values for these permutation tests were therefore generated by fitting a beta cumulative distribution with two free parameters (a and b) to the permutation distribution of the statistic and comparing the observed statistic to the fitted beta distribution. Results of fitting the beta distribution are shown in supplementary figures 4B & 5B.

All imaging analyses included sex, parent-reported race and ethnicity, household income, parental education and scanner as categorical fixed covariates and age as a standardised (z-scored) continuous fixed covariate. For vertex-wise cortical thickness analyses, standardised global mean thickness was included as a covariate, and for vertex-wise cortical surface area analyses, standardised total surface area was included. Analyses were conducted in Matlab R2017a. Details of the empirical simulations used to validate the MOSTest procedure in this context and test the false positive rate and sensitivity of the MOSTest compared to the min-P can be found in the supplementary methods.

Although previous literature as has suggested that mean CTH and total CSA are independent, it is not clear whether the regionalisation of CTH would be independent of total CSA. Indeed, controlling for mean CTH does not control for brain size in the same way as total CSA. Therefore, we repeated the MOSTest for relative CTH adding total CSA as an additional covariate.

#### Empirical Simulations

In order to validate the false positive rate of the MOSTest method and directly compare the sensitivity of the MOSTest and the standard approach we used empirical simulations. We simulated the vertex-wise imaging data as a null or non-null signal plus noise. The noise was estimated by calculating the residuals from regressing the covariates of no interest only on the imaging data and permuting the subject labels. For the non-null case, we simulated the vertex-wise imaging data using empirical estimated effect sizes. This ensured the distribution and signal-to-noise ratio of the simulated data was similar to our empirical associations. For this we used the estimates of *β* from the association between the NIH picture sequence task and relative CTH. For the null case, the estimates of *β* were set to zero. These data were simulated for 100 random samples at different sample sizes (N=100, 500, 1000, 2000, 3000, 4000, 5000). For each random sample, the association between the behaviour of interest (NIH picture sequence) and the null and non-null simulated data were assessed using 1000 permutations of both the MOSTest statistic and min-P. For the MOSTest, the covariance matrix used in all iterations was estimated from the null distribution of mass univariate Wald statistics from 10,000 permutations of the full sample.

For each sample size and statistic, we calculated sensitivity as the proportion of significant permutation tests for the non-null simulated data (true positives) and specificity as 1 minus the proportion of significant permutation tests for the null simulated data (false positives) over the 100 iterations.

### Statistical Analysis 2: quantifying the magnitude of the association between brain structure and cognition using a Bayesian PVS (PVS_B_)

The Bayesian-PVS (PVS_B_) was calculated for each cognitive task to determine the behavioural variance explained by the vertex-wise cortical morphology. All behavioural and imaging data were pre-residualised using the covariates of no interest prior to calculation of the PVS_B_. For the imaging data, the global CSA and CTH measures specific to each modality were also included in this pre-residulation. This allowed us to determine the unique association between relative cortical morphology and cognition and compare this to the predictive power of brain structure without controlling for global measures. The method used here is outlined in details by Zhao and colleagues (16). The association between each imaging phenotype and each cognitive task was modelled using the mass univariate approach in which the behaviour of interest was modelled independently at each vertex using a general linear model (GLM). A Bayesian posterior mean effect size was then calculated at each vertex to take into account the correlation amongst vertices. This method was adapted from the LDpred framework originally developed to improve accuracy for polygenic risk scores (44). The PVS predicting behaviour from cortical morphometry was then computed for each subject as the linear-weighted sum of the estimated Bayesian effects and the pre-residualized cortical morphometry vector. This measure thus harnesses the explanatory power of all of the vertices with respect to behaviour. The computed PVS is then compared with the observed neurocognitive assessment in order to provide an estimate for how much variance in the observed behaviour can be predicted using the vertex-wise imaging phenotype.

In order to generate an unbiased PVS_B_ for every subject we used a leave-one-out 10 fold cross-validation procedure. The Bayesian parameters were estimated in the training set (90% of full sample) and multiplied with the imaging phenotype of participants in the test set (10% of full sample). This was repeated 10 times for each fold until a PVS_B_ was calculated for every participant in the full sample. The subjects in each fold were randomly selected based on unique family IDs, such that subjects within the same family were always within the same fold. The association between the imaging phenotype and behaviour across the whole sample was calculated as the squared correlation (R^2^) between the observed behaviour and the predicted behaviour (the PVS_B_). This process was repeated for four imaging phenotypes: CSA, relative CSA (controlling for total CSA), CTH. relative CTH (controlling for mean CTH). This procedure provides a conservative estimate of the variance explained, when applied to a finite sample, due to errors in the vertexwise effect size estimates.

In order to determine the unique variance in cognitive performance predicted by relative CTH independent of total CSA, we calculated another PVS_B_ for relative CTH including total CSA as a covariate in the pre-residualisation of the imaging data for each of the cognitive tasks.

## DATA AND CODE AVAILABILITY

All ABCD data is openly available to download from the NIMH Data Archive. All MATLAB scripts for the analyses conducted in this study are available on both the ABCD GitHub, https://github.com/ABCD-STUDY, and the Precimed GitHub https://github.com/precimed.

## ACKNOWLEDGEMENTS

The authors wish to thank Anderson Winkler for his helpful comments in the implementation of the permutation scheme used for the MOSTest. We’d also like to thank the youth and families participating in the Adolescent Brain Cognitive Development (ABCD) Study and all ABCD staff. Data used in the preparation of this article were obtained from the Adolescent Brain Cognitive Development (ABCD) Study (https://abcdstudy.org), held in the NIMH Data Archive (NDA). This is a multisite, longitudinal study designed to recruit more than 10,000 children age 9-10 and follow them over 10 years into early adulthood. The ABCD Study is supported by the National Institutes of Health and additional federal partners under award numbers U01DA041022, U01DA041028, U01DA041048, U01DA041089, U01DA041106, U01DA041117, U01DA041120, U01DA041134, U01DA041148, U01DA041156, U01DA041174, U24DA041123, U24DA041147, U01DA041093, and U01DA041025. A full list of supporters is available at https://abcdstudy.org/federal-partners.html. A listing of participating sites and a complete listing of the study investigators can be found at https://abcdstudy.org/Consortium_Members.pdf. ABCD consortium investigators designed and implemented the study and/or provided data but did not all necessarily participate in analysis or writing of this report. This manuscript reflects the views of the authors and may not reflect the opinions or views of the NIH or ABCD consortium investigators. The ABCD data repository grows and changes over time. The data was downloaded from the NIMH Data Archive ABCD Collection Release 2.0.1 (DOI: 10.15154/1504041).

