## Supplementary Material for "Determining the association between regionalisation of cortical morphology and cognition in 10,145 children"

SUPPLEMENTARY TABLES AND FIGURES

| **Type of Variable** | **Variable** | **NDA/DEAP Variable Name** | **Instrument** |
| --- | --- | --- | --- |
| *Covariates of no interest* | Age | interview_age | Developmental history questionnaire (dhx01) |
|  | Sex | sex | Developmental history questionnaire (dhx01) |
|  | Self-declared race/ethnicity | race.4level**  hisp** | Parent Demographics Survey (pdem02) – computed DEAP variable |
|  | Income | household.income | Parent Demographics Survey (pdem02) – computed DEAP variable |
|  | Parental education | high.educ | Parent Demographics Survey (pdem02) – computed DEAP variable |
|  | Scanner ID | mri_info_deviceserialnumber | MRI Info (abcd_mri01) |
| *Family ID* | Family | rel_family_id | ACS Post Stratification Weights (acspsw02) |
| *MRI QC* | Freesurfer QC | fsqc_qc | FreeSurfer QC (freesqc01) |
| *Cognitive Measures* | Toolbox Oral Reading Recognition | nihtbx_reading_uncorrected | Youth NIH TB Summary Scores (abcd_tbss01) |
|  | Toolbox Picture Vocabulary Task | nihtbx_picvocab_uncorrected | Youth NIH TB Summary Scores (abcd_tbss01) |
|  | Toolbox Flanker Task | nihtbx_flanker_uncorrected | Youth NIH TB Summary Scores (abcd_tbss01) |
|  | Toolbox Dimensional Card Sorting Task | nihtbx_cardsort_uncorrected | Youth NIH TB Summary Scores (abcd_tbss01) |
|  | Toolbox Pattern Speed Task | nihtbx_pattern_uncorrected | Youth NIH TB Summary Scores (abcd_tbss01) |
|  | Toolbox Picture Sequence Task | nihtbx_picture_uncorrected | Youth NIH TB Summary Scores (abcd_tbss01) |
|  | Toolbox List Working Memory Task | nihtbx_list_uncorrected | Youth NIH TB Summary Scores (abcd_tbss01) |
|  | Toolbox Fluid Composite Score | nihtbx_fluidcomp_uncorrected | Youth NIH TB Summary Scores (abcd_tbss01) |
|  | Toolbox Crystallised Composite Score | nihtbx_cryst_uncorrected | Youth NIH TB Summary Scores (abcd_tbss01) |
|  | Toolbox Total Composite Score | nihtbx_totalcomp_uncorrected | Youth NIH TB Summary Scores (abcd_tbss01) |
|  | Matrix reasoning | pea_wiscv_trs | Pearson Scores (abcd_ps01) |
|  | Little Man Task | lmt_scr_efficiency | Little Man Task Summary Scores (lmtp201) |
|  | RAVLT | pea_ravlt_sd_trial_i_tc+pea_ravlt_sd_trial_ii_tc+pea_ravlt_sd_trial_iii_tc+pea_ravlt_sd_trial_iv_tc+pea_ravlt_sd_trial_v_tc | Pearson Scores (abcd_ps01) |

**Supplementary Table 1.** List of all variables used in this study with corresponding variable names and the instruments in which they can be found in the ABCD data dictionary and data release. Only participants with complete data on all of these variables, available T1-weighted MRI scans and whose imaging data passed the freesurfer QC flag (fsqc_qc=1) were included in the analyses.** *We computed race/ethnicity by changing the race of all those who endorsed being Hispanic to ‘Hispanic’ to create a 5-level race/ethnicity variable as seen in release 1.1 and 2.0.*

|  | **Analysed sample** |
| --- | --- |
| **n** | 10145 |
| **Age, in months (mean (sd))** | 119.02 (7.48) |
| **Sex = M (%)** | 5288 (52.1) |
| **Parental education (%)** |  |
| < HS Diploma | 460 ( 4.5) |
| HS Diploma/GED | 942 ( 9.3) |
| Some College | 2624 (25.9) |
| Bachelor | 2611 (25.7) |
| Post Graduate Degree | 3508 (34.6) |
| **Household income (%)** |  |
| [<50K] | 2841 (28.0) |
| [>=50K & <100K] | 3155 (31.1) |
| [>=100K] | 4149 (40.9) |
| **Race/ethnicity (%)** |  |
| White | 5411 (53.3) |
| Black | 1484 (14.6) |
| Asian | 211 ( 2.1) |
| Other | 989 ( 9.7) |
| Hispanic | 2050 (20.2) |
| **Scanner (%)** |  |
| HASH03db707f | 423 ( 4.2) |
| HASH11ad4ed5 | 480 ( 4.7) |
| HASH1314a204 | 506 ( 5.0) |
| HASH311170b9 | 339 ( 3.3) |
| HASH31ce566d | 43 ( 0.4) |
| HASH3935c89e | 959 ( 9.5) |
| HASH48f7cbc3 | 29 ( 0.3) |
| HASH4b0b8b05 | 387 ( 3.8) |
| HASH4d1ed7b1 | 352 ( 3.5) |
| HASH5ac2b20b | 462 ( 4.6) |
| HASH5b0cf1bb | 559 ( 5.5) |
| HASH5b2fcf80 | 255 ( 2.5) |
| HASH65b39280 | 307 ( 3.0) |
| HASH69f406fa | 130 ( 1.3) |
| HASH6b4422a7 | 318 ( 3.1) |
| HASH7911780b | 358 ( 3.5) |
| HASH7f91147d | 86 ( 0.8) |
| HASH96a0c182 | 535 ( 5.3) |
| HASHa3e45734 | 291 ( 2.9) |
| HASHb640a1b8 | 471 ( 4.6) |
| HASHc3bf3d9c | 439 ( 4.3) |
| HASHc9398971 | 199 ( 2.0) |
| HASHd422be27 | 419 ( 4.1) |
| HASHd7cb4c6d | 497 ( 4.9) |
| HASHdb2589d4 | 495 ( 4.9) |
| HASHe3ce02d3 | 86 ( 0.8) |
| HASHe4f6957a | 505 ( 5.0) |
| HASHe76e6d72 | 15 ( 0.1) |
| HASHfeb7e81a | 200 ( 2.0) |

**Supplementary Table 2. Descriptive characteristics of the analysed sample of n=10,145 participants.** This table shows the numbers and percentage (in brackets) of participants within each of the levels of the categorical demographic variables (sex, parental education, household income, race and ethnicity and scanner). For the continuous measure of age the mean and standard deviation across the sample is shown.

|  | **CSA** | | **CTH** | | **CTH (controlling for total CSA)** | |
| --- | --- | --- | --- | --- | --- | --- |
|  | $\boldsymbol{z}_{\boldsymbol{MINP}}^{\boldsymbol{2}}$ | $\boldsymbol{\chi}_{\boldsymbol{MOST}}^{\boldsymbol{2}}$ | $\boldsymbol{z}_{\boldsymbol{MINP}}^{\boldsymbol{2}}$ | $\boldsymbol{\chi}_{\boldsymbol{MOST}}^{\boldsymbol{2}}$ | $\boldsymbol{z}_{\boldsymbol{MINP}}^{\boldsymbol{2}}$ | $\boldsymbol{\chi}_{\boldsymbol{MOST}}^{\boldsymbol{2}}$ |
| Toolbox Oral Reading Recognition Task | 1.23E-03 | 5.03E-07 | 1.48E-04 | 8.56E-13 | 1.14E-02 | 1.32E-07 |
| Toolbox Picture Vocabulary Task | 1.80E-03 | 8.16E-09 | 1.08E-02 | 3.30E-09 | 7.30E-02 | 6.90E-06 |
| Toolbox Flanker Task | 3.40E-02 | 8.70E-04 | 1.50E-02 | 9.56E-06 | 4.37E-02 | 8.09E-04 |
| Toolbox Dimensional Card Sorting Task | 3.23E-03 | 3.43E-04 | 4.15E-03 | 9.04E-06 | 9.23E-03 | 9.60E-08 |
| Toolbox Pattern Processing Speed Test | 3.18E-02 | 1.37E-05 | 4.01E-02 | 7.09E-08 | 2.87E-02 | 8.19E-06 |
| Toolbox Picture Sequence Memory Test | 8.64E-02 | 2.10E-04 | 3.69E-03 | 8.34E-08 | 4.10E-03 | 1.14E-11 |
| Toolbox List Working Memory Test | 2.70E-03 | 4.60E-06 | 5.01E-07 | 1.54E-11 | 9.89E-03 | 3.78E-17 |
| **Toolbox Fluid Composite Score** | 9.95E-04 | 3.99E-11 | 1.17E-04 | 2.39E-22 | 1.08E-03 | 1.08E-24 |
| **Toolbox Crystallised Composite Score** | 9.72E-05 | 3.07E-09 | 2.60E-05 | 5.94E-18 | 2.19E-01 | 4.38E-09 |
| **Toolbox Total Composite Score** | 1.65E-03 | 1.25E-10 | 2.09E-05 | 1.06E-29 | 4.06E-03 | 1.74E-25 |
| Matrix Reasoning | 2.24E-03 | 1.87E-04 | 3.05E-04 | 5.50E-10 | 1.19E-03 | 1.86E-06 |
| Rey-Auditory Verbal Learning Task (RAVLT) | 5.75E-03 | 8.81E-05 | 8.02E-03 | 2.29E-04 | 3.42E-02 | 3.94E-05 |
| Little Man Task (LMT) | 1.34E-01 | 1.73E-07 | 1.78E-02 | 1.08E-06 | 3.03E-02 | 1.00E-08 |

**Supplementary Table 3.** Theoretical p-values for the association between cortical morphology and each of the cognitive measures for the standard approach ($P_{MIN})$ and the MOSTest ($\chi_{MOST}^{2}$). The far right columns show the associations results for relative CTH predicting cognition controlling for total CSA. Across all of these analyses, more of the cognitive measures showed significant associations with cortical morphology using the MOSTest compared to the standard approach. The composite scores are highlighted in bold. All significant associations according to the permutation test are highlighted in red.

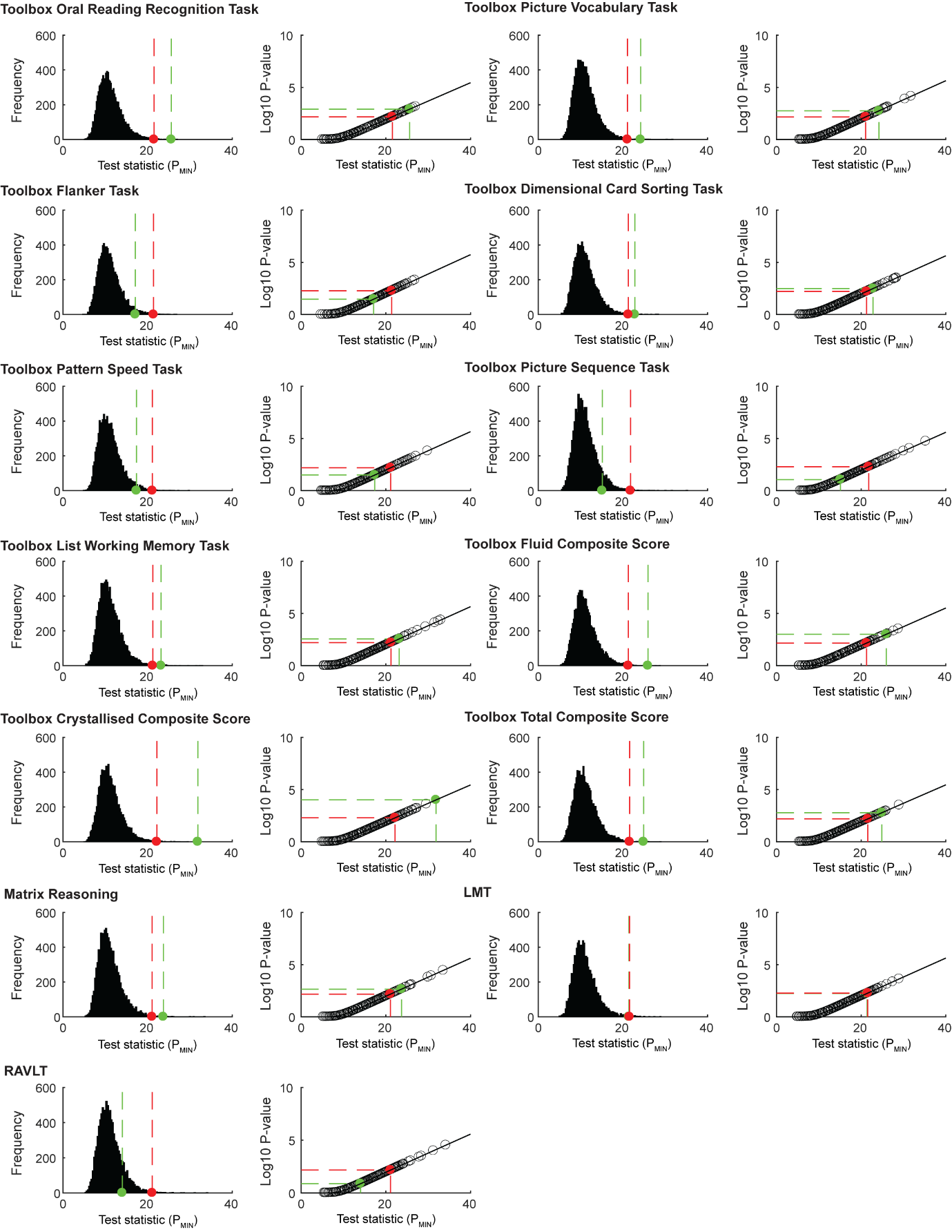

**Supplementary Figure 1. Standard approach omnibus permutation test for each cognitive task and relative CSA.** The most significant vertex ($P_{MIN}$) was used as the observed statistic to determine an association between the brain and behaviour. All of the left side plots show the permuted null distribution of this statistic for each cognitive task. The observed, non-permuted test statistic is shown in green and the Bonferroni corrected alpha level in red. All of the right side plots show a beta cumulative density function fit to the null distribution in order to extrapolate a theoretical p-value for the observed statistic.

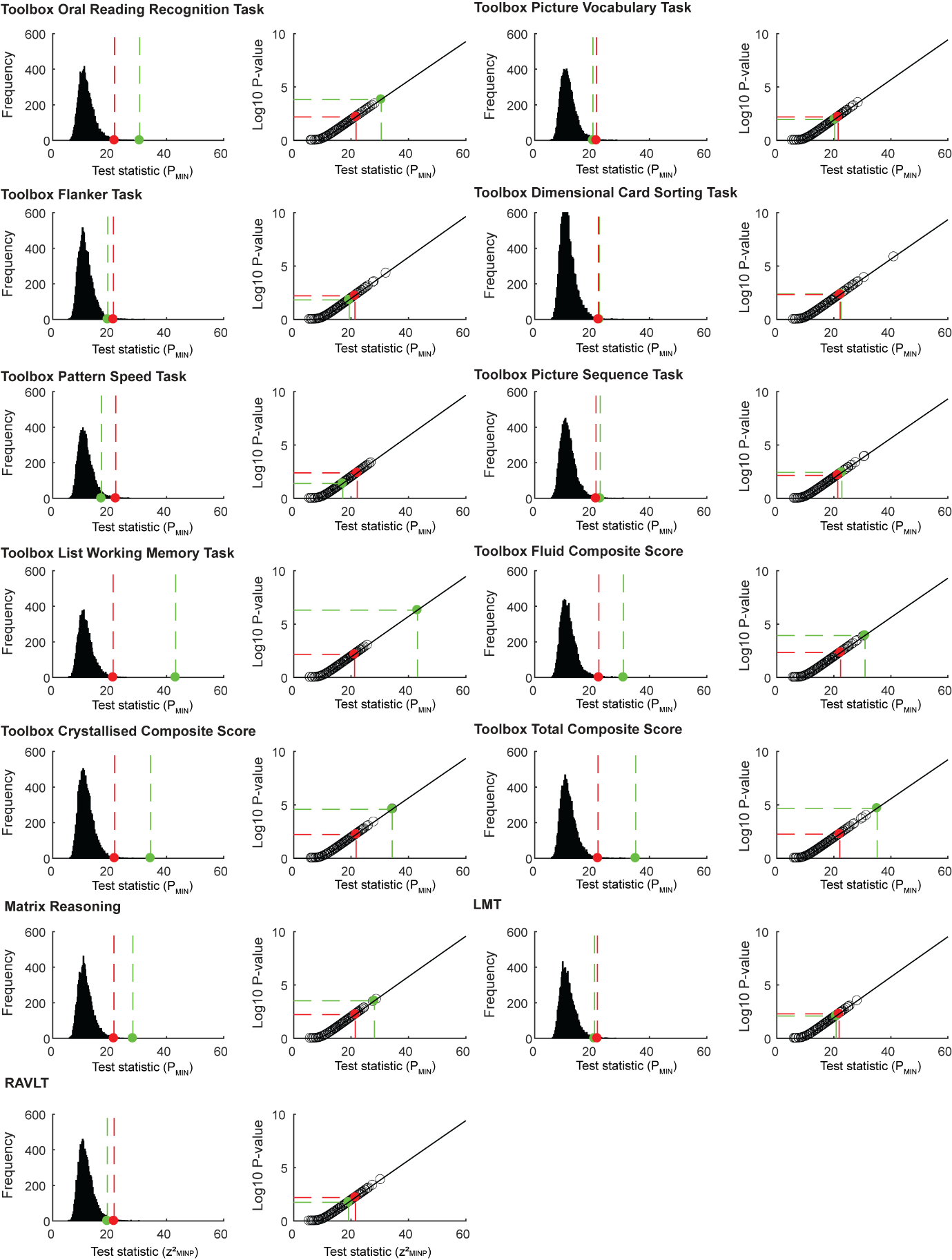

**Supplementary Figure 2. Standard approach omnibus permutation test for each cognitive task and relative CTH.** The most significant vertex ($P_{MIN}$) was used as the observed statistic to determine an association between the brain and behaviour. All of the left side plots show the permuted null distribution of this statistic for each cognitive task. The observed, non-permuted test statistic is shown in green and the Bonferroni corrected alpha level in red. All of the right side plots show a beta cumulative density function fit to the null distribution in order to extrapolate a theoretical p-value for the observed statistic.

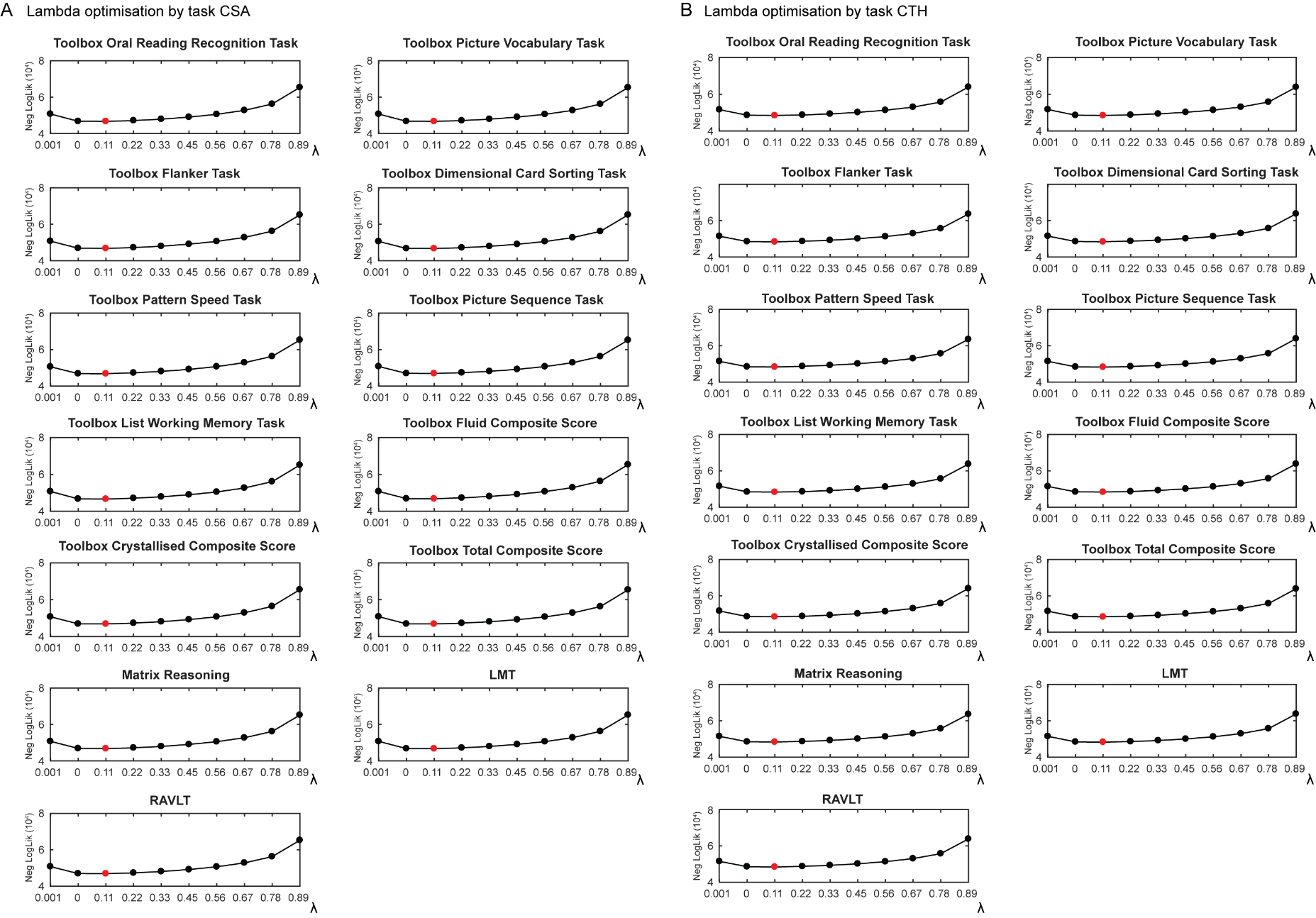

**Supplementary Figure 3. Lambda optimisation for the regularisation of the estimated covariance structure of the imaging data.** The calculation of the multivariate statistic, the Mahalanobis distance, involves a regularised covariance matrix estimated from the matrix of permuted z-statistics. The magnitude of regularisation is determined by the penalty factor, λ (Eq 3 in the main text). Here we optimise λ based on the goodness of fit of the permuted distribution and the gamma CDF for each of the cognitive measures for (A) CSA and (B) CTH for different values of λ. In all cases the best fit was found for a λ value of 0.1. This was used in all analyses in this paper.

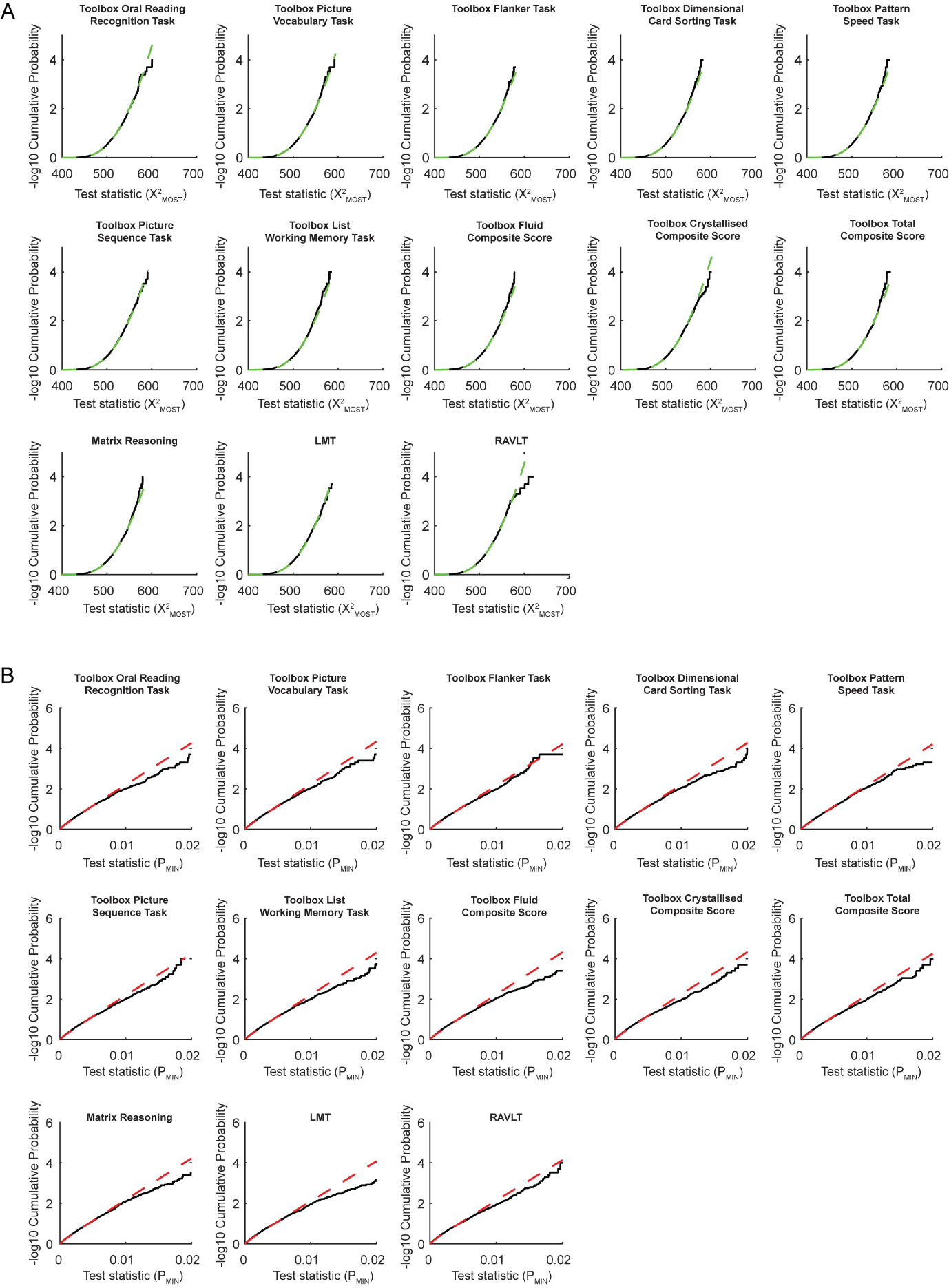

**Supplementary Figure 4. Cumulative density functions (CDF) for permuted and fitted data for CSA associations.** These plots show the CDF of the permuted test statistic (black) and the predicted CDF after fitting either a gamma distribution (green dotted) to the permuted multivariate statistic ($\chi_{MOST}^{2}$; A) or beta distribution (red dotted) to the permuted univariate statistic ($P_{MIN}$; B) for each cognitive task. The fitted CDF was extrapolated to provide an approximated p-value for the observed statistic.

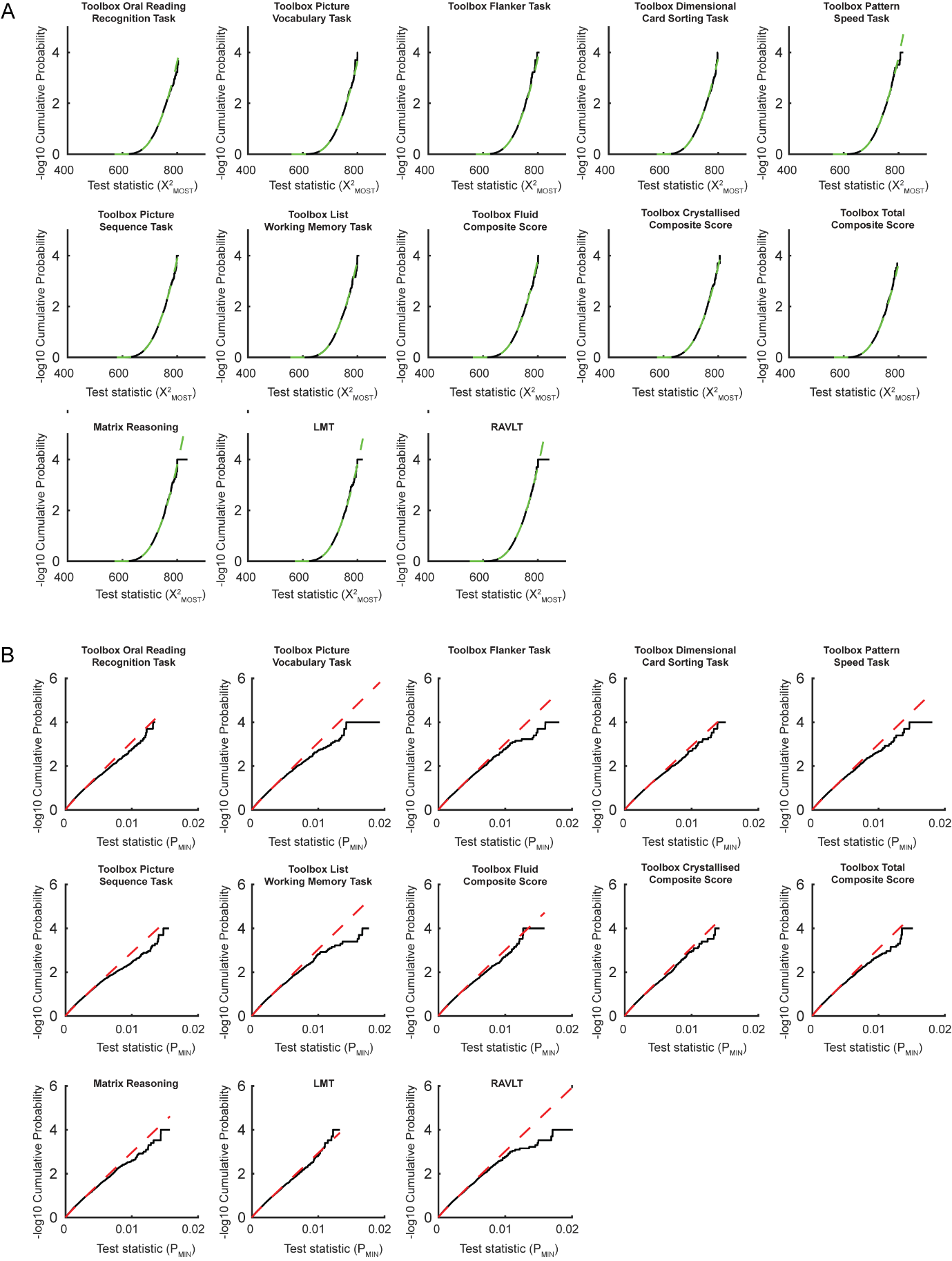

**Supplementary Figure 5. Cumulative density functions for permuted and fitted data for CTH associations.** These plots show the CDF of the permuted test statistic (black) and the predicted CDF after fitting either a gamma distribution (green dotted) to the permuted multivariate statistic ($\chi_{MOST}^{2}$; A) or beta distribution (red dotted) to the permuted univariate statistic ($P_{MIN}$; B) for each cognitive task. The fitted CDF was extrapolated to provide an approximated p-value for the observed statistic.

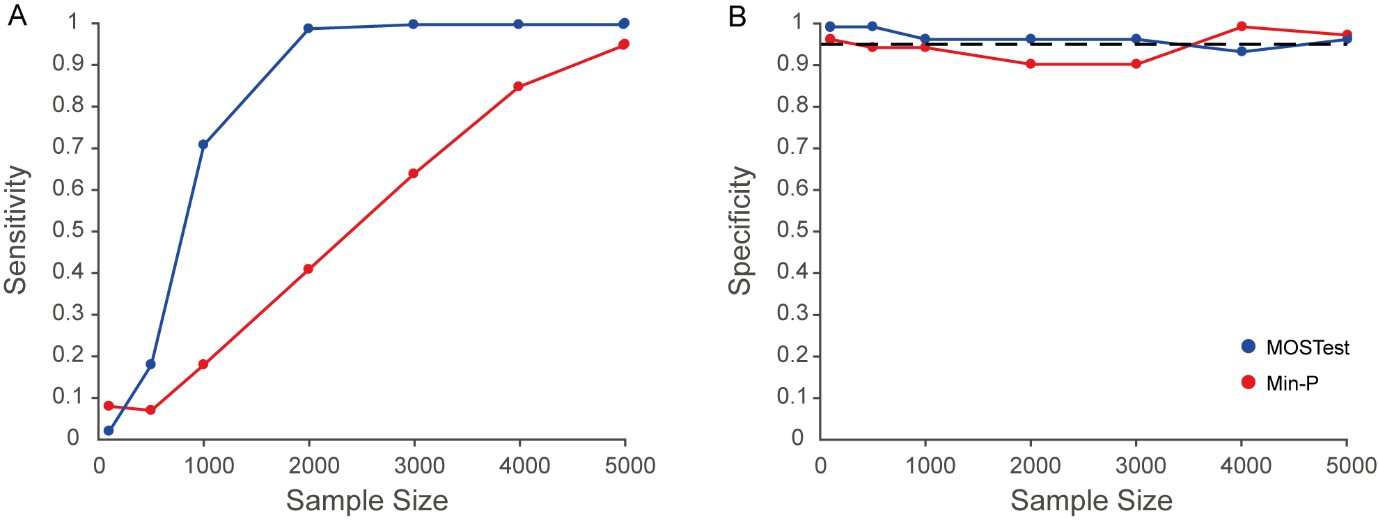

**Supplementary Figure 6. Detection performance of the MOSTest and the standard approach using empirical simulations.** Here we show the sensitivity (A) and specificity (B) of the two omnibus tests, MOSTest (blue) and min-P (red), across 100 iterations of random samples at different sample sizes (N=100, 500, 1000, 2000, 3000, 4000, 5000). For a given sample size, sensitivity was calculated as the proportion of significant permutation tests across 100 random samples for the association between the *non-null* simulated vertex-wise imaging data and the behaviour of interest. Specificity was calculated as 1 - the proportion of significant permutation tests across 100 iterations for the association between the *null* simulated vertex-wise imaging data and the behaviour of interest. The MOSTest showed increased sensitivity for detection compared to the standard approach and maintained a false positive rate around 5%. The false positive rate for the standard approach, however, increased to around 10% for some of the simulations.

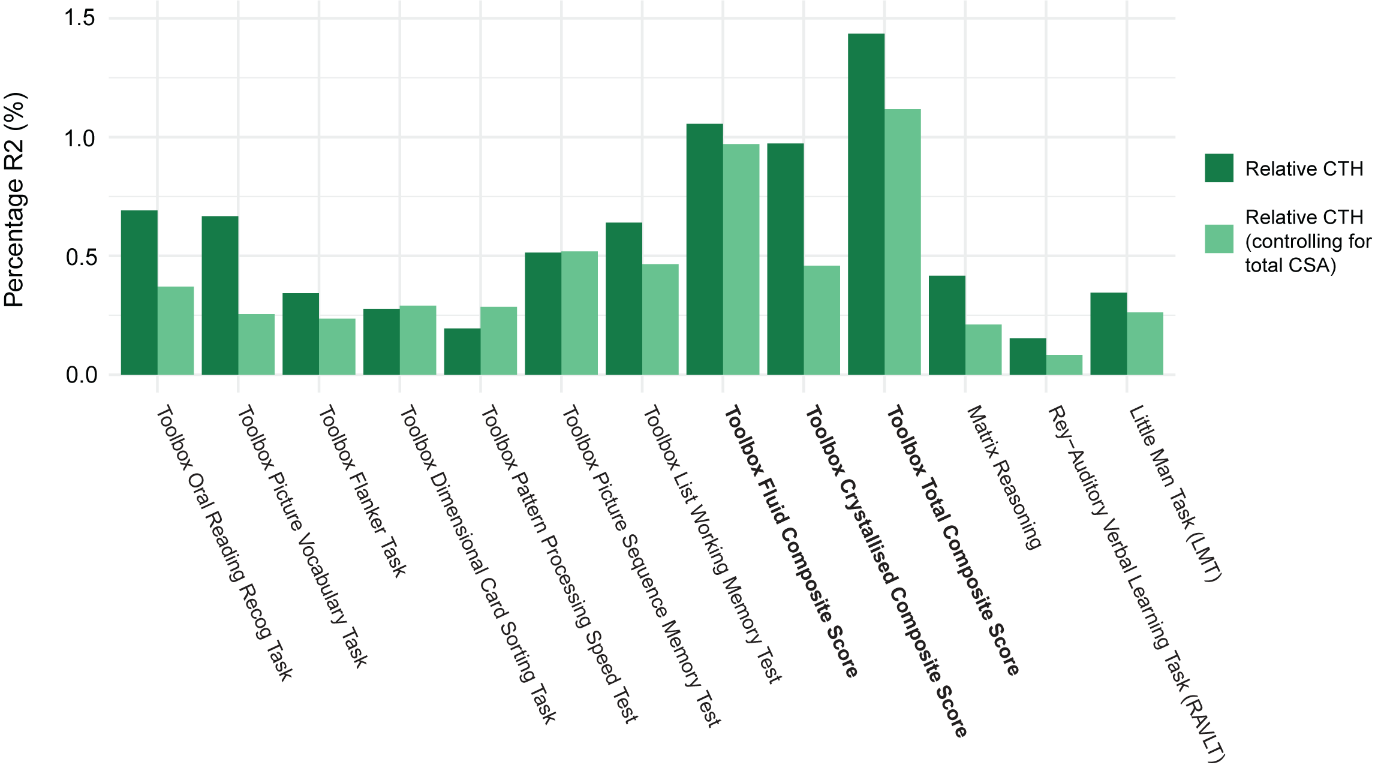

**Supplementary Figure 7. Partial mediation of total CSA on the regionalisation of CTH predicting cognition.** This figure shows the variance in cognitive performance (percentage R^2^) explained by relative CTH (dark green; only controlling for mean CTH) and relative CTH after controlling for both mean CTH and total CSA (light green) in order to determine the unique variance associated with relative CTH independent of total CSA. The crystallised measures showed a greater reduction in R^2^ associated with relative CTH after controlling for total CSA compared to the fluid measures.
